# Scaffolding of long read assemblies using long range contact information

**DOI:** 10.1101/083964

**Authors:** Jay Ghurye, Mihai Pop, Sergey Koren, Chen-Shan Chin

## Abstract

**Motivation:** Long read technologies have made a revolution in *de novo* genome assembly by generating contigs of size orders of magnitude more than that of short read assemblies. Although the assembly contiguity has increased, it still does not span a chromosome or an arm of the chromosome, resulting in an unfinished chromosome level assembly. To address this problem, we develop a scalable and computationally efficient scaffolding method that can boost the contiguity of the assembly by a large extent using genome wide chromatin interaction data such as Hi-C. Particularly, we demonstrate an algorithm that uses Hi-C data for longer-range scaffolding of *de novo* long read genome assemblies.

**Results:** We tested our methods on two long read assemblies of different organisms. We compared our method with previously developed method and show that our approach performs better in terms of accuracy of scaffolding.

**Availability:** The software is available for free use and can be downloaded from here: https://github.com/machinegun/hi-c-scaffold

## 1 Introduction

The advent of massively parallel sequencing technologies has made generation of billions of reads possible at a very low cost per sequenced base. Despite the progress made in de novo assembly algorithms, the quality of short read assemblies is far from the quality necessary for effective further analysis due to fundamental limit- the read length is shorter than repeat lengths for the majority of repeat classes (Nagarajan and Pop, 2009; Bresler *et al.*, 2013). For example, a short read *de novo* assemblies of human genome are highly fragmented compared to the chromosomes of the *H.sapien* reference (Gnerre *et al.*, 2011; Li *et al.*, 2010). Thus, high throughput sequencing technology has reached a point where increasing the number of short reads does not significantly improve assembly quality.

Recent advances in single-molecule sequencing technologies have provided reads almost 100 times longer than second generation methods (Schatz *et al.*, 2010). Most prominently, Pacific Biosciences’ single molecule real time (SMRT^®^) sequencing delivers reads of lengths up to 50 Kbp (Eid *et al.*, 2009) whereas Oxford Nanopore’s nanopore sequencing can deliver read lengths greater than 10 Kbp (Lee *et al.*, 2014). Such read lengths drastically reduce the complexity caused by repeats during the assembly process. However, these long reads suffer from low accuracy which requires new algorithms for assembly. Despite the higher error rates, SMRT sequencing has random error model (Koren *et al.*, 2012; Ono *et al.*, 2013) due to which near perfect assembly is possible (Lam *et al.*, 2014). Hence by sampling the genome at sufficient coverage, SMRT sequencing has been used to produce assemblies of unprecedented continuity (Koren *et al.*, 2013; Chin *et al.*, 2013; Ribeiro *et al.*, 2012; Berlin *et al.*, 2015).

Various strategies have been explored to increase the continuity of *de novo* genome assemblies. Some of these strategies are end sequencing of fosmid clones (Gnerre *et al.*, 2011), fosmid clone dilution pool sequencing (Chinwalla *et al.*, 2002), optical mapping (Schwartz *et al.*, 1993; Dong *et al.*, 2013; Shelton *et al.*, 2015; English *et al.*, 2015), linked-read sequencing (Zheng *et al.*, 2016; Zook *et al.*, 2016) and synthetic long reads (McCoy *et al.*, 2014; Koren and Phillippy, 2015; Madoui *et al.*, 2015). Some of the newer technologies like Hi-C use proximity ligation and massively parallel sequencing to probe the three-dimensional structure of chromosomes within the nucleus, with interacting regions captured by pair-ended reads (Lieberman-Aiden *et al.*, 2009; Duan *et al.*, 2010). In the data generated by Hi-C protocol, the probability of intrachromosomal contacts is on an average much higher than the that of interchromosomal contacts (Burton *et al.*, 2013; Kaplan and Dekker, 2013). Another important property is that the probability of interaction decays rapidly with increasing genomic distance, even regions separated by several hundred megabases on the same chromosome are more likely to interact than the regions on different chromosomes (Lieberman-Aiden *et al.*, 2009).

In this work, we make use of the genome-wide chromatin interaction data sets generated by the Hi-C protocol to linearly arrange the pre-assembled contigs along entire chromosomes. We develop a scaffolding tool SALSA (Simple AssembLy ScAffolder) based on a computational method that exploits the information of genomic proximity in Hi-C data sets for long range scaffolding of *de novo* genome assemblies. We tested SALSA on human and goat genome assembly. Our method can produce centromere to telomere scaffolds of chromosomes in most cases and telomere to telomere scaffolds in best cases.

## 2 Related Work

Several efforts have been made to use Hi-C data to scaffold the ‘draft stage’ short read assemblies. (Burton *et al.*, 2013) developed a computational approached in their tool LACHESIS that combined Hi-C data with pair ended sequencing data to generate chromosome level scaffolds. They used their methods to scaffold *de novo* assemblies of human, mouse and *Drosophilia Melanogaster.* LACHESIS uses the alignment of Hi-C reads to contigs to clusters contigs into 1 cluster per chromosome with hierarchical clustering. It then orients and orders the contigs in each cluster to obtain final scaffolds. One of the drawbacks of this method is that it needs the number of clusters to be pre-specified. This can not be applied to scaffolding contigs of genomes when the number of chromosomes in the organisms are unknown. (Kaplan and Dekker, 2013) developed a method for scaffolding based on statistical techniques. Their method uses the hierarchical clustering method similar to LACHESIS, but it predicts the number of clusters by itself. The major drawback of their method is that they do not orient the contigs in each cluster, thereby not providing complete information needed for scaffolding. Their method also works with an assumption that all contigs are of the same size, which does not hold true in the case of long read assemblies since there can be contigs which are tens of Mb long whereas there can be shorter contigs generated by the assembler. Since both of these methods rely on hierarchical clustering, it is expensive to compute all vs all link scores for all the contigs thereby causing the scalability issues. Another drawback of both methods is that they do not provide the way to detect and correct misassemblies generated by the assembler. If such errors are not corrected, then that would result in erroneous scaffolds in the end and may also propagate errors across multiple scaffolds causing misjoins.

In our work, we address the issues in the previous methods. Our method does not need the number of clusters to be pre-specified. We also provide an option to detect certain types of misassemblies in contigs before scaffolding them.

## 3 Methods

### 3.1 Aligning Hi-C Reads

Hi-C reads were aligned to SMRT read assemblies using BWA (Li and Durbin, 2009) with default parameters. Reads with mapping quality < 30, which included the reads mapped more than once were removed from further analysis. Also, only the read pairs with both reads in the pair aligned to contigs are considered for scaffolding.

### 3.2 Detection of Mis-assemblies

Contigs obtained from assemblers may contain mis-assemblies (Phillippy *et al.*, 2008). We provide a method to detect and correct these misassemblies similar to the one described in (Putnam *et al.*, 2016) using the mapping of Hi-C data to the assembled contigs. For each read pair, it’s physical coverage is defined as the total bases spanned by the sequence of reads and the gap between the two reads when mapped to contigs (Figure 1). We also define, per base physical coverage for each base in the contig as the number of read pairs’ physical coverage it is part of. Using these definitions, we compute the physical coverage for each base of all the contigs in the assembly. The misassembly can be detected by the sudden drop in per-base physical coverage in a contig. A particular threshold below which if per base physical coverage falls for contiguous regions in the genome, we call it a misassembly and break contigs at that point. To do this efficiently, we use a variation of Kadane’s algorithm for maximum sum subarray problem (Simon and Kadane, 1975). We find the subarray in the array of physical coverage where coverage is consistently low compared to the adjacent regions and use that as the signal for misassembly.

**Fig. 1.**
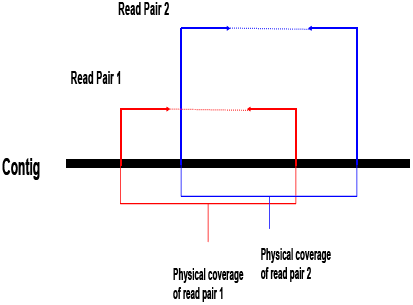
Physical coverage induced by Hi-C read pair. The solid arrows denote the read pair and dotted line denote the gap between the reads in the pair.

### 3.3 Graph Construction and Link Scoring

We use an idea similar to the string graph formulation in (Myers, 2005) to construct the scaffold graph. The scaffold graph *G*(*V*, *E*, *W*) consists of nodes *V* which represent the end of contigs, edgs *E* representing the edges implied by Hi-C read pairs between ends of two contigs and weight function *W* to assign weight to each edge. The ends of each contig are annotated by two tags, *B* and *E* where *B* stands for the beginning of the contig and *E* stands for the end of the contig. Using this concept of node, there are 4 types of edges in the graph, *BE* joining beginning of first contig to end of second, *EB* joining end of the first contig to beginning of second, *BB* joining beginning of the first contig to beginning of second contig and *EE* joining end of the first contig to end of second (Figure 2). Using raw counts of Hi-C read pairs shared between ends of two contigs is not the correct way to score the edges because of length biases since longer contigs with a large genomic distance between them tend to share more read pairs compared to two short or one long and one short contig with much lesser genomic distance between them. To address this issue, we define an edge weight function in such a way that it reduces such length biases. We define a length cutoff *l* and consider the read pairs mapped in the region of length *l* at both ends (*B* and *E*) of contigs. Our normalization of the edge weight is based on how many times the restriction enzyme used in Hi-C protocol cuts the region of length *l* and divide the counts of read pairs by this number. Putting all this together, the edge weight function is expressed as:

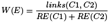

where *C*1 and *C*2 are the contigs yielding the edge, *links*(*C*1, *C*2) is the number of hi-c links present in region of length *l* at the end of contigs and *RE*(*C*1) and *RE*(*C*2) is the number of sites cut by the restriction enzyme in region *l* at the end of *C*1 and *C*2. This gives us the normalized edge weights which we use for scaffolding.

**Fig. 2.**
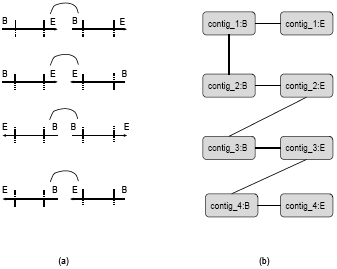
(a) Type of edges present in the scaffold graph. We consider Hi-C read pairs originating from the end of contigs. Since a contig can be reverse complemented, there are 4 possible links, (b) Scaffold Graph *G*. There are two nodes in *G* corresponding to each contig. There is always an edge between ’B’ node and ’E’ nodes of same contig.

Once we calculate the edge weights, we construct graph *G* as follows. We first sort all the edges in decreasing order of their weights given by *W*. After this, we remove all the edges which have very low number of read pairs shared between them which account for sequencing errors. Once edges are sorted and filtered, we construct *G* according to Algorithm 1. We greedily add edges to *G* only if both the nodes corresponding to the edge are not present in *G*. In the end, we add edges between *B* and *E* ends of same contigs to *G* which completes the graph construction. *G* can contain some cycles due to the following reason: we can add an edge between both *B* and *E* ends of two contigs (*BB* and *EE* edges) and later add edges between *B* and *E* ends of same contigs. However, this kind of cycle can easily be removed by removing the lower cost edge among *BB* and *EE* edges. Once we remove cycles from *G*, we get final scaffold graph which we use for further analysis (Figure 2).

### 3.4 Scaffold Construction

Before explaining the scaffold construction algorithm, we prove following lemmas to understand the properties of *G*.

#### Lemma 3.1.

*G has no nodes with degree greater than 2.*

#### Algorithm 1 Scaffold Graph Construction

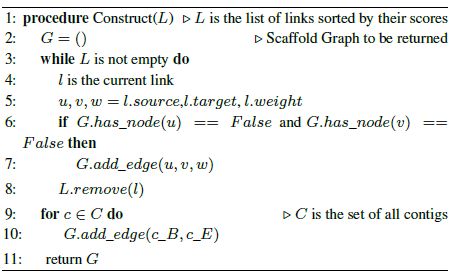

#### Proof

While constructing *G*, we add edges at most twice for each node. First when we have no edge associated to that node and second when we add an edge between *B* and *E* ends of the contig associated with that node. If some node has degree greater than 2, it would mean that we added an edge to that node apart from the cases described previously, which is a contradiction.

#### Lemma 3.2.

*Each connected component of G has exactly two nodes of degree 1.*

#### Proof

We know from the construction of *G* that *G* has no cycles. We can prove this for some connected component *C* of *G* and the argument can be applied to all connected components. Since *G* has no cycles, *C* will have at least one node with degree 1. In the first case, suppose *C* Chas exactly one node of degree 1. This implies that we have at least two edges originating from all other nodes in *C*. It would mean that there exists at least one node in *C* with a degree at least 3. This is a contradiction because of lemma 3.1. In the second case, suppose *C* has more than two nodes of degree 1. It would mean that there exists at least one node in *C* with degree 3. This is again a contradiction due to lemma 3.1.

Knowing these properties about *G* we construct scaffolds as described in Algorithm 2. First, a threshold *th* is decided for a scaffold to qualify as a seed scaffold. If a scaffold has a number of contigs greater than *th*, it is marked as seed scaffold. For each connected component of *G*, we first find out two nodes *u* and *v* with degree 1. We will always find such nodes due to lemma 3.2. After this, the path connecting *u* and *v* is found in the connecting component. Since all the nodes in the connected component have degree either 1 or 2, there will always be just 1 path connecting *u* and *v*. If this path has the number of contigs greater than *th*, we mark this path as seed scaffold, otherwise, it is marked as the small scaffold.

Even after edge weight normalization, there can still be some length biases resulting in the omission of smaller contigs from the seed scaffolds. To account for this, we develop a method to insert the contigs in small scaffolds into seed scaffolds. First, for each contig in small scaffolds, exactly one seed scaffold is assigned to it based on the total edge weight of the edges in the original scaffold graph connecting this contig to all the contigs in seed scaffold. After this, each contig is tested for insertion into its corresponding seed scaffolds in both the orientations at all possible position. It is inserted at the position where it maximizes the total weight of the scaffold. Once all the contigs are inserted into seed scaffolds, it leaves us with the final scaffolds. The algorithm is sketched in Algorithm 2.

## 4 Dataset

For NA 12878, a human genome used in 1000 genomes project (Siva, 2008), we used the assembly generated at Icahn School of Medicine (Genebank assembly accession GCA_001013985.1). This assembly was performed using Celera Assembler (Koren *et al.*, 2012) and had 21235 contigs with N50 of 1.55 Mb. For the scaffolding of NA 12878 assembly, Hi-C data produced from human ESCs (hESCs) (Dixon *et al.*, 2012) was used. The hESC replicates 1 and 2 were used (NCBI SRA Accession: GSM862723, GSM892306), consisting of total 734M read-pairs. For *Capra hircus* genome we used the data presented in (Bickhart *et al.*, 2016). *Capra hircus* is a domestic goat of San Clemente breed. The assembly performed using long reads with CA PBcR pipeline had contig NG50 of 4.15 Mb. These contigs were scaffolded using Irys optical mapping data, resulting in increased scaffold size. All the sequencing data and intermediate assemblies are publically available^1^. We used Hi-C data (NCBI SRA Accession: SRX1910977) with both contig assembly and Irys optical map scaffolds to perform scaffolding and compared the extended scaffolds with the scaffolds given by LACHESIS.

### Algorithm 2 Scaffold Construction

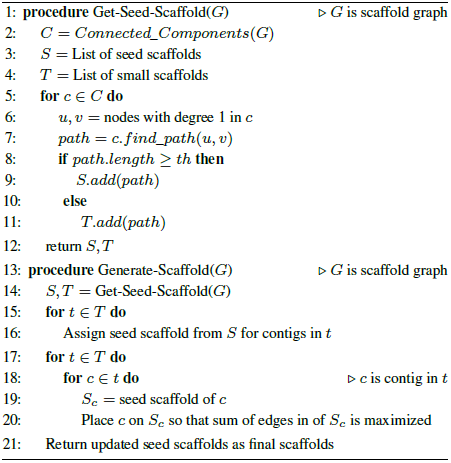

## 5 Results

### 5.1 Contact Probability of Hi-C data

We aligned Hi-C reads for NA12878 to GRCh38 human reference using BWA mem (Li and Durbin, 2009) with default parameters. If both mates in the read pair are aligned to the same chromosome, it provides an evidence that Hi-C data gives valuable information about the intrachromosomal contact. For each chromosome, we count how many read pair have both mates mapped to that chromosome and how many reads have just one of the mates mapped to that chromosome. Using this information, we compute the probability of intrachromosomal and interchromosomal contact for each chromosome. It can be seen from Figure 3 that the probability of intrachromosomal contact is much higher than that of interchromosomal contact.

**Fig. 3.**
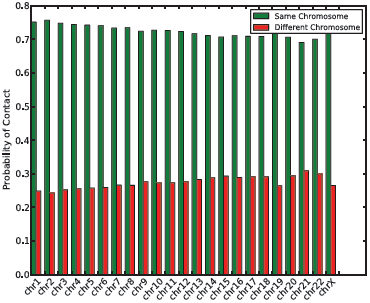
The probability of contact calculated based on read mapping to GRCh38 reference

To further understand the contact probability at contig level, we aligned reads to NA12878 contigs. We also aligned contigs to GRCh38 reference using Mummer (Delcher *et al.*, 1999). For each contig that mapped to chromosome 1, we calculated its average interaction frequency to contigs belonging to the all other chromosomes. Figure 4 shows the box plot of the distribution of average interaction frequency of all contigs of chromosome 1 among themselves and with the contigs in all other chromosomes. We found that the average interaction frequency strongly separates inter from intra-chromosomal interactions.

**Fig. 4.**
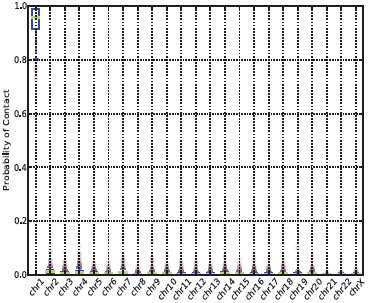
The probability of contact of all the contigs in chromosome 1 to all other contigs.

### 5.2 Scaffolding assemblies of two genomes

We tested the effectiveness of our approach for the chromosome scale *de novo* assembly of two genomes. We started with two assemblies, one for NA 12878 with contig N50 of 1.55 Mb and the other for goat genome with contig N50 of 4.15 Mb. After aligning Hi-C read pairs to these assemblies, we used our algorithm to construct the scaffold graph and later to orient and order contigs. For NA12878 assembly, it gave 1555 scaffolds with NG50 of 60.02 Mb. For *Capra hircus* assembly, it gave 127 scaffolds with NG50 of 68.64 Mb. Table 1 shows the statistics of the contigs and scaffolds.

**Table 1.** Contig and Scaffold statistics for NA 12878 and Capra hirrus

| Metric | NA12878 | *Capra hircus* |
| --- | --- | --- |
| **Number of Contigs** | 21235 | 33767 |
| **Contig N50** | 1.55 Mb | 3.86 Mb |
| **Number of Scaffolds** | 1555 | 127 |
| **Scaffold NG50** | 60.02 Mb | 58.64 Mb |
| **Total Bases** | 2.77 Gb | 2.94 Gb |

### 5.3 Comparison with LACHESIS

To evaluate the quality of the scaffolds, we aligned NA12878 scaffolds to human GRCh38 reference. For goat scaffolds, we aligned them to recently published goat reference genome (BioProject PRJNA290100) (Bickhart *et al.*, 2016) using *nucmer* program (parameters: -c 1000) in MUMmer package (Delcher *et al.*, 1999). The quality of alignments was assessed using *dnadiff* program (Phillippy *et al.*, 2008), which evaluates the draft assemblies by comparing with the reference genome based on a set of metrics. We particularly focus on four metrics. First one is a number of breakpoints which is defined as the number of alignments which are not end to end with reference to a particular scaffold. A second metric is a number of relocations which is the number of breaks in the alignment where adjacent aligned sequences are in the same sequence but not the correct order. This accounts of ordering errors in the scaffold construction. A third metric is a number of translocations, which counts the number of breaks in the alignment where adjacent sequences are in different chromosomes. This accounts for the inter-chromosomal join errors in the scaffold. A last metric is a number of inversions, which is the number of breaks in the alignment where adjacent sequences are inverted with respect to each other. This accounst for the orientation errors caused due to scaffolding algorithm.

Table 2 shows the comparison of scaffolds generated by SALSA and LACHESIS for NA12878 assembly. For the scaffolds generated by SALSA, 80.82% bases could align to reference whereas only 40.70% bases from the scaffolds generated by LACHESIS could align to the reference. Although the number of bases aligned to the reference is significantly low in LACHESIS scaffolds, the relocation, translocation and inversion errors are much higher in LACHESIS compared to SALSA. Particularly, the number of relocations is significantly high in LACHESIS implying that LACHESIS performs a lot of errors in ordering contigs belonging to a particular chromosome (Bickhart *et al.*, 2016).

**Table 2.** Evaluation of scaffolds generated by SALSA and LACHESIS for the human NA12878 assembly

| Metric | SALSA | LACHESIS |
| --- | --- | --- |
| Number of Scaffolds | 1555 | 23 |
| Total Bases | 2.92 Gb | 2.79 Gb |
| % Aligned Bases | 80.82% | 40.70% |
| Breakpoints | 14010 | 6820 |
| Relocations | 19 | 76 |
| Translocations | 4 | 9 |
| Inversions | 42 | 65 |

Table 3 shows the comparison of scaffolds generated by SALSA and LACHESIS for *Capra hircus* assembly. In this case, both the scaffolds have an almost similar percent of aligned bases to reference. Even in this case, LACHESIS performs a lot of orientation and ordering errors compared to SALSA. SALSA produces 67 and 105 orientation and ordering errors in the scaffolds, which is significantly lesser than 374 and 439 orientation and ordering errors produced by LACHESIS. In addition to this, SALSA produces lesser inter-chromosomal joins(213) compared to LACHESIS(601).

**Table 3.** Evaluation of scaffolds generated by SALSA and LACHESIS for the Capra hircus assembly

| Metric | SALSA | LACHESIS |
| --- | --- | --- |
| Number of Scaffolds | 127 | 990 |
| Total Bases | 2.44 Gb | 2.62 Gb |
| % Aligned Bases | 99.88% | 99.85% |
| Breakpoints | 8514 | 14035 |
| Relocations | 67 | 374 |
| Translocations | 213 | 601 |
| Inversions | 105 | 439 |

Figure 5 and Figure 6 show the alignment dotplot for human and goat scaffolds respectively when aligned to respective reference genomes. In the case of NA12878 scaffolds, it can be seen that (Figure 5 A) the scaffolds generated by LACHESIS are very fragmented and lack continuity. The scaffolds generated by SALSA (Figure 5 B) are continuous, contain lesser orientation and ordering errors and much more consistent with the reference than LACHESIS scaffolds. Alignment dot plots for each chromosome can be found in the supplementary material (Supplementary Figure S1). In the case of goat scaffolds, it can be seen that although LACHESIS produces continuous scaffolds, it incurs a lot of large scale orientation and ordering errors (Figure 6 A). In contrast, SALSA is able to produce the continuous scaffolds with lesser amount of orientation and ordering errors compared to LACHESIS thereby producing scaffolds that are consistent with the reference to a large extent.

**Fig. 5.**
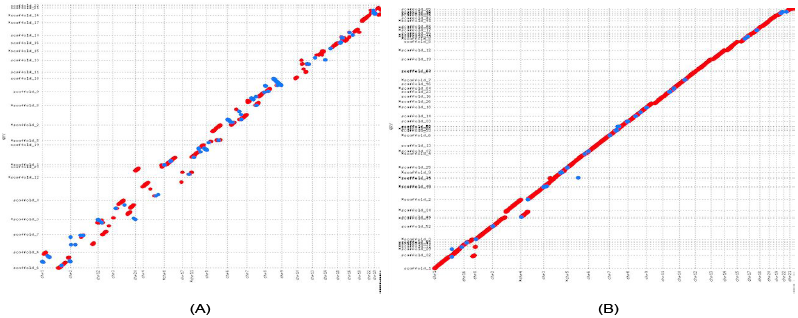
(A) The alignment dotplot of Lachesis scaffolds tor NA 12878. (B) The alignment dotplot for NA12878 scaffolds generated by SALSA.

**Fig. 6.**
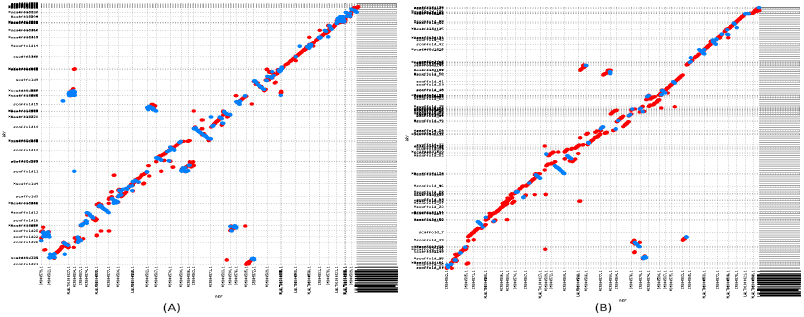
(A) The alignment dotplot of Lachesis scaffolds for Capra hircus. (B) The alignment dotplot for Captra hircus scaffolds generated by SALSA.

### 5.4 Scaffolding Optical Map Scaffolds

(Bickhart *et al.*, 2016) also made scaffolds of goat assembly generated using optical map data available publically. We used these scaffolds as a starting point for our scaffolding method and did further scaffolding using Hi-C data. The initial scaffold assembly had 1575 scaffolds with N50 of 23.08 Mb. After scaffolding Hi-C data with SALSA, it gave 90 scaffolds, and scaffolding with LACHESIS gave 596 scaffolds. We evaluated these scaffold using the metrics described before and the results are shown in Table 4. It can be observed that even though the initial N50 of the input assembly is high, LACHESIS is still prone to ordering errors compared to SALSA. However, in this case, SALSA produced 3.17% more orientation errors than LACHESIS. The alignment dotplots for both the scaffolds are shown in supplementary material (Supplementary Figure S2).

**Table 4.** Evaluation of scaffolds generated by SALSA and LACHESIS for Capra hircus assembly generated using optical map data

| Metric | SALSA | LACHESIS |
| --- | --- | --- |
| Number of Scaffolds | 90 | 596 |
| Total Bases | 2.22 Gb | 2.61 Gb |
| NG50 | 46.64 Mb | 87.34 Mb |
| % Aligned Bases | 97.00% | 99.91% |
| Breakpoints | 10718 | 14741 |
| Relocations | 118 | 167 |
| Translocations | 185 | 407 |
| Inversions | 130 | 126 |

## 6 Conclusion and Discussion

In this work, we show that genome-wide interaction data sets like Hi-C are a good source of information to scaffold the pre-assembled contigs into scaffolds. Since we use long read assemblies at the start which mostly span through highly repetitive regions in the genome, we do not need to do repeat masking as previous methods and hence derive more continuous scaffolds. We derive a weight function to normalize the scores of Hi-C links which reduces the length biases caused by long contigs. Since long read assemblies have non-uniform contig lengths, edge weight normalization plays an important role in deriving correct scaffolds. We provide a method to correct misassemblies in the starting assembly so that these errors do not propagate through the further scaffolding process. We used string graph-based approach along with some greedy heuristics to derive scaffolds. We tested our methods on assemblies of two different organisms namely human and goat. Our method showed significant improvements on the same dataset over previously used method. Since our method does not need the number of chromosomes to be specified apriori, it can be applied for scaffolding the assemblies of organisms where we don’t know the number of chromosomes beforehand. However, designing a clustering method that clusters the contigs without knowing the actual number of desired clusters is needed to estimate the number of chromosomes in an unknown organism. To avoid errors in clustering, we can first orient and order the contigs with respect to each other and cluster the resulting scaffolds as we would have larger and more continuous parts of the genome to cluster compared to contigs which are smaller portions of the genome. Most of the orientation and ordering errors in our method were in the repetitive regions near centromeres and telomeres. One potential solution to overcome this problem is to do all pairwise alignment of contigs and trim the contigs so that these repetitive regions are masked. However, there is a serious computational bottleneck associated to this.

There are still several open questions. Our method needs some parameter tuning to get good scaffolds. We plan to incorporate optimal parameter detection at runtime so that the onus of parameter tuning is taken away from the user of the tool. There are other chromatin interaction datasets like Dovetail Chicago libraries (Putnam *et al.*, 2016) developed recently. We plan to extend our method so that it becomes generic to all such datasets and adapts to their chromosomal contact model.

## Acknowledgements

We thank Ashton Trey Belew for helping with running LACHESIS tool for comparative analysis. We also thank Chris M. Hill for helpful discussions.

## Funding

Sergey Koren was supported by the Intramural Research Program of the National Human Genome Research Institute, National Institutes of Health. This study utilized the computational resources of the Biowulf system at the National Institutes of Health, Bethesda, MD (http://biowulf.nih.gov).

## Competing Financial Interest

Chen-Shan Chin is an employee and shareholder of Pacific Biosciences, a company commercializing DNA sequencing technologies.

## Footnotes

1 Assemblies and sequencing data information can be found here: https://gembox.cbcb.umd.edu/goat/index.html

